# Substitution spectrum and selection at G-quadruplexes in great ape telomere-to-telomere genomes

**DOI:** 10.1101/2025.08.24.671323

**Authors:** Xinru Zhang, Saswat Mohanty, Francesca Chiaromonte, Kateryna D. Makova

**Author notes:** Corresponding author: Correspondence to Kateryna D. Makova.

## Abstract

G-quadruplexes (G4s) are noncanonical DNA secondary structures formed by runs of guanines (stems) connected by other nucleotides (loops). These structures are enriched at regulatory regions such as promoters, CpG islands, untranslated regions (UTRs), enhancers, and replication origins, where they play key roles in transcription and replication. Although prior studies have demonstrated that G4s exhibit higher mutation rates than canonical DNA, little is known about the substitution patterns and selection acting specifically on G4 stems and loops. In this study, we utilized Telomere-to-Telomere (T2T) genome assemblies from human and two non-human great apes (chimpanzee and Bornean orangutan) to analyze substitution spectra and selective constraints within G4s, focusing on differences between stems and loops. We observed that fixed nucleotide substitutions leading to the gain or loss of G4 structures are more frequently located at stems, while those in G4s conserved across species are more often found at loops. On the other hand, single nucleotide polymorphisms had higher frequencies at stems than loops for all G4s, with a particularly high difference for singleton polymorphisms, suggesting higher mutation rates at stems than loops. To evaluate selection, we employed two approaches: we computed the ratio of substitution to polymorphism frequencies at stems vs. loops and performed phylogenetic modeling using PhyloFit. Both methods consistently revealed that stems of shared G4s experience stronger purifying selection than loops, particularly at promoters, enhancers, and UTRs. Our results provide novel insights into the sequence variation and selection of G4s, informing our understanding of their contributions to genome evolution and function.

**Significance Statement:** G-quadruplexes (G4s) are non-canonical DNA structures that influence transcription, genome stability, and epigenetic regulation, yet their evolutionary dynamics in primates remain poorly understood. Leveraging recent T2T genome assemblies, we conducted a sequence-level analysis of G4 evolution across three ape lineages and used two methods to infer selection in G4. Shared G4s and species-specific G4s display distinct evolutionary signatures. Using PhyloFit to estimate substitution rates and the Hudson–Kreitman–Aguadé test to contrast divergence with polymorphism, we found stems under markedly stronger purifying selection than loops, especially within promoters, CpG islands, and 5′UTRs. This pattern indicates that maintaining stem integrity is functionally critical and evolutionarily conserved. Our findings reveal how selective constraints vary both within G4 motifs and across genomic landscapes, offering insights for future studies on their functional importance and structural stability.

## Introduction

G-quadruplexes (G4s) are alternative (non-B) DNA secondary structures forming guanine-rich regions (Sen and Gilbert 1988; Sundquist and Klug 1989). G4s are composed of stems and loops (Fig. 1A); stems are tracts of at least three guanines and loops, which can contain any nucleotides, connect neighboring stems. A canonical G4 has four stems and three loops. Four guanines (one on each stem) form a square planar arrangement, linked by Hoogsteen hydrogen bonds. Since each stem typically contains three guanines, a canonical G4 forms three planar arrangements (Varshney et al., 2020; Spiegel, Adhikari, and Balasubramanian, 2020; Bochman, Paeschke, and Zakian,(Varshney et al. 2020; Spiegel, Adhikari, and Balasubramanian 2020; Bochman, Paeschke, and Zakian 2012). G4 structures usually form at specific sequence motifs, e.g., G_x_N_1-12_G_x_N_1-12_G_x_N_1-12_G_x_, where x ≥ 3 and N can be any nucleotide (later called ‘G4 motifs’). There are also instances where G4 structures form at sequences that do not adhere to such G4 motifs. For example, the stacking of two planar arrangements can also form G4 structures (Zhang et al. 2010; Guiblet et al. 2018).

**Fig. 1.**
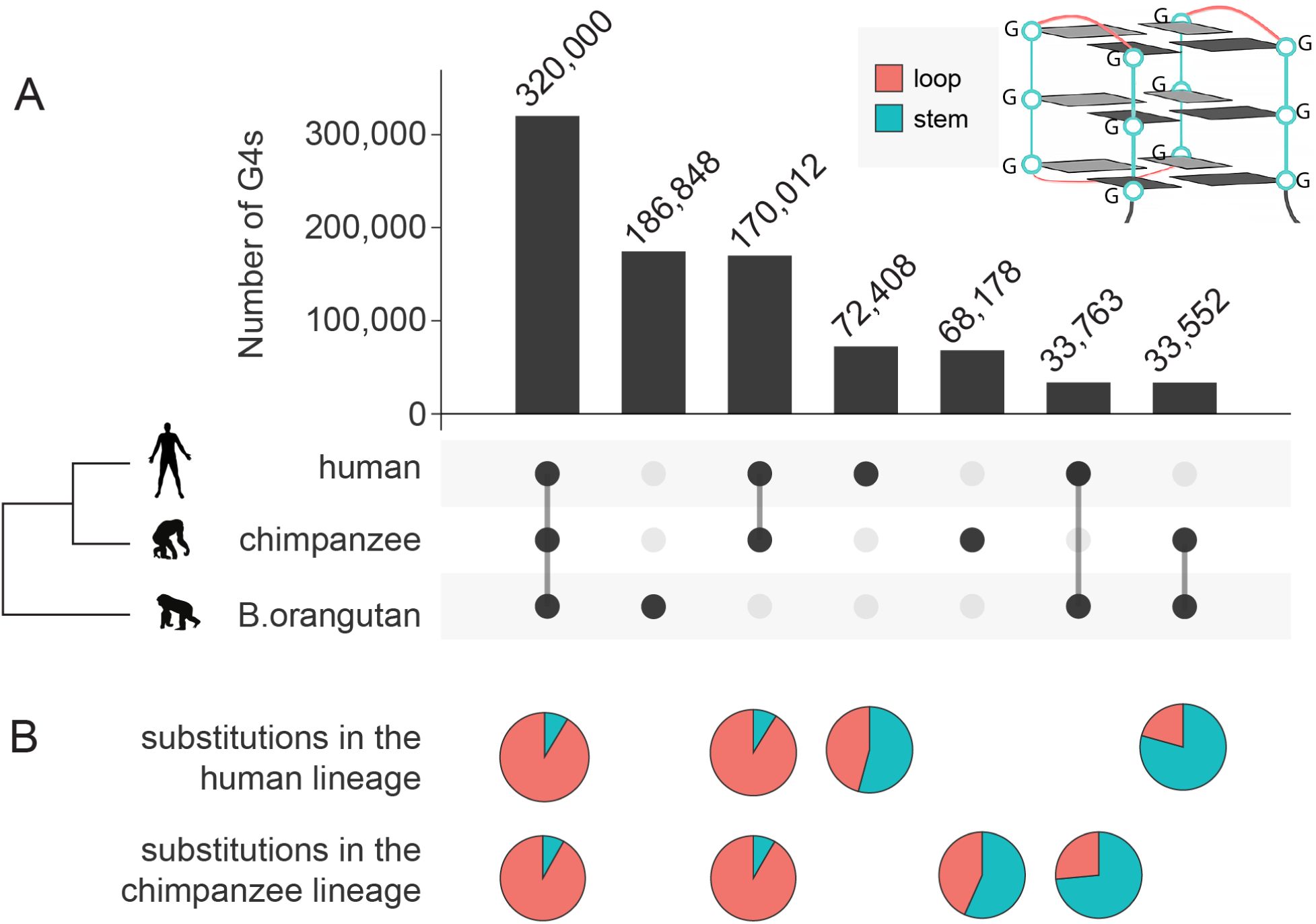
Evolutionary groups of G4s and nucleotide substitution distribution in stems vs. loops. (**A**) The interspecific sharing of Quadron-predicted G4s at orthologous locations of the three great ape T2T genomes. Animal symbols are from phylopic (Michael Keesey, n.d.). G4s were classified into seven evolutionary groups based on their presence and absence in three species. The schematic representation of a typical G4 structure is shown in the upper right corner. (**B**) Proportions of substitutions in stems (blue) vs. loops (red) within G4s. The rows correspond to G4 evolutionary groups. The directions of substitutions were inferred using parsimony.

G4 motifs are not randomly distributed across the genome; they show uneven densities along the chromosomes. In particular, they are enriched at promoters, upstream of transcription start sites, downstream of 3’ untranslated regions (UTRs), and at telomeres, and they are depleted at exons (Guiblet, DeGiorgio, et al. 2021; Mohanty, Chiaromonte, and Makova 2025). An enrichment of G4 motifs was also found at exon-intron boundaries (Georgakopoulos-Soares et al. 2022), CpG islands, 5’UTRs, and replication origins (Guiblet et al. 2018; Mohanty, Chiaromonte, and Makova 2025). Extensive studies have revealed multiple biological functions of G4 structures in chromatin organization, transcription, replication, and recombination (reviewed in (Wang and Vasquez 2023)).

In a recent study on genetic divergence and diversity in G4s and other types of non-B DNA (Guiblet, Cremona, et al., 2021), we found that G4s exhibited higher diversity and divergence compared to B DNA, suggesting their greater mutability. Additionally, we discovered that stable G4s had higher mutability than unstable ones. Another recent study comparing selective pressures at G4s and non-G4 regions found purifying selection favoring motif retention and sequence conservation at functional G4 loci (Guiblet, DeGiorgio, et al. 2021). For example, G4s at promoters, CpG islands, and UTRs were found to be subject to purifying selection, whereas G4s at the nontranscribed strands of exons evolved neutrally. However, whether substitution spectra and selective pressures differ within G4 motifs, particularly between stems and loops, is a question that remains unanswered to date.

To address these questions, we leveraged the newly released gapless and complete T2T genome assemblies for humans (Nurk et al. 2022; Miga et al. 2020; Rhie et al. 2023) and great apes (Yoo et al. 2024; Makova et al. 2024). These assemblies, which resolved challenging genomic regions, provide an opportunity for a more detailed and complete analysis of G4 evolution. We started with an assumption that selection ought to be stronger at stems than loops of functional G4s, and we inferred selective constraints at stems and loops of G4s using two approaches: (1) comparing substitution rates with polymorphism rates at stems *vs.* loops; (2) fitting phylogenetic models separately to stem and loop alignments using PhyloFit (Hubisz, Pollard, and Siepel 2011).

## Results

### Classification of G4s based on evolutionary history

Using Quadron (Sahakyan et al. 2017), we annotated G4s in the T2T genomes of human, chimpanzee, and Bornean orangutan (later called ‘B. orangutan’). This resulted in 740,644 G4s in the human genome (Nurk et al. 2022; Rhie et al. 2023), 752,616 G4s in the chimpanzee genome (Makova et al. 2024; Yoo et al. 2024), and 738,918 G4s in the B. orangutan genome (Makova et al. 2024)).

We then extracted the sequences orthologous to human G4s from the chimpanzee and B. orangutan T2T genomes using pruned six-way CACTUS (Paten et al. 2011) alignments of whole-genome assemblies of six great ape species (bonobo, chimpanzee, human, gorilla, B. orangutan, and Sumatran orangutan). A total of 591,862 human G4s were found in these six-way alignments. Based on CACTUS alignments and considering the three focal species of interest (human, chimpanzee, and B. orangutan), human G4s were divided into four groups (Table S1, Fig. 1): (1) shared with chimpanzee and B. orangutan (‘ape G4s’), (2) shared with chimpanzee only (‘Hominini G4s’), (3) shared with B. orangutan only (‘G4s lost in chimpanzee’), and (4) present in human only (‘human-specific G4s’).

Similarly, we extracted the sequences of G4s orthologous to chimpanzee G4s from the human and B. orangutan genomes, using the same six-way pruned CACTUS alignments. A total of 591,742 G4s were extracted with this procedure. They were similarly divided into four groups: (1) shared with human and B. orangutan (identical to the ‘ape G4s’ group above), (2) shared with human (identical to the ‘Hominini G4s’ group above), (3) shared with B. orangutan (‘G4s lost in human’), and (4) present in chimpanzee only (‘chimpanzee-specific G4s’).

### Substitution spectrum at G4s

We next analyzed divergence between human and chimpanzee, and between human and B. orangutan, at G4s annotated in human and/or chimpanzee genomes. We applied a parsimony-based approach to reconstruct the ancestral sequence and infer the presence or absence of G4s in the common ancestor of these species (Table S1). The distribution of substitutions in the human and chimpanzee lineages is shown in Fig. 2 and Table S2.

**Fig. 2.**
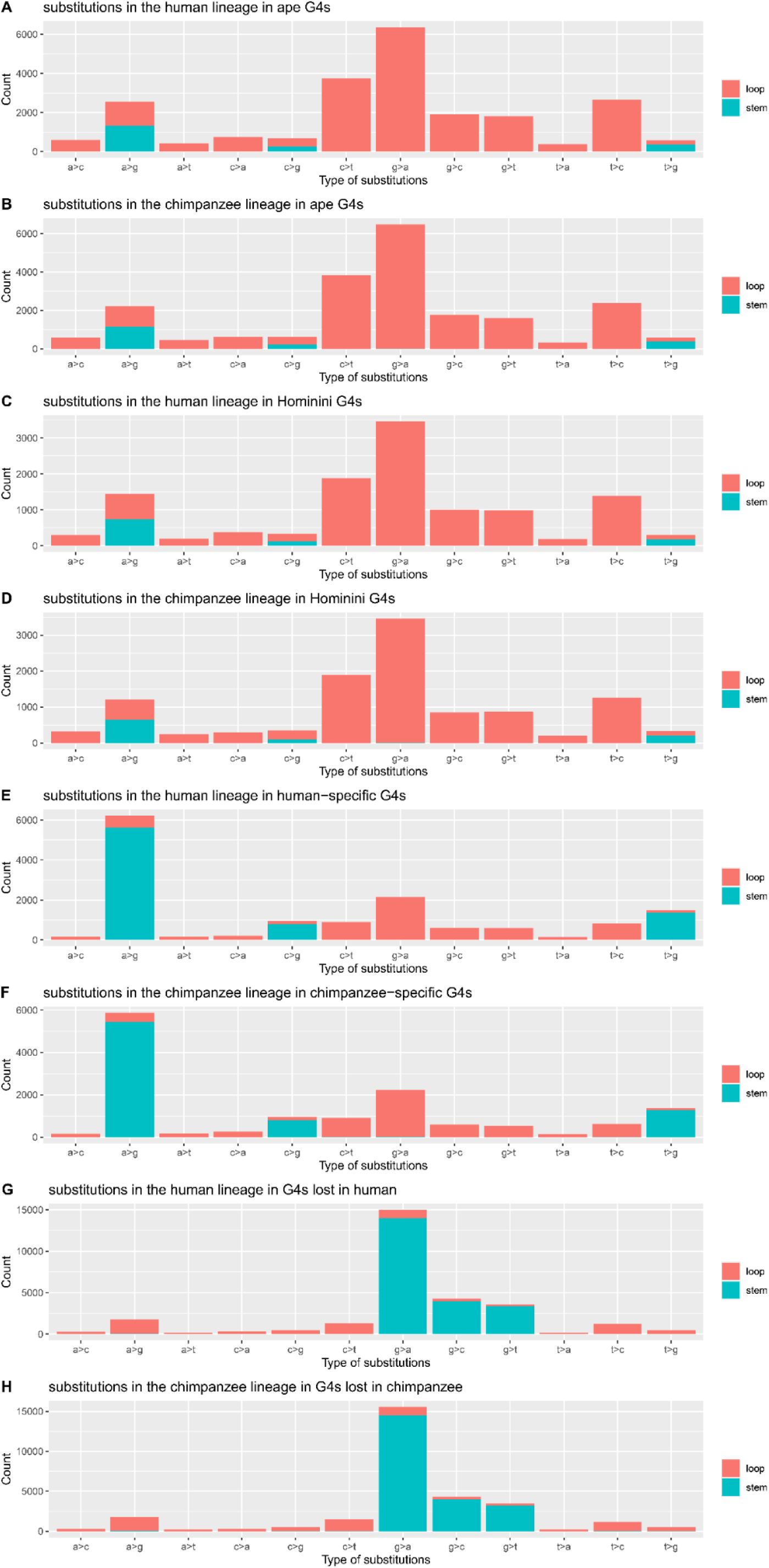
Nucleotide substitution spectra of G4s. Substitutions in stem and loop sequences are shown in blue and red, respectively. Distribution of nucleotide substitutions (**A**) from the human-chimpanzee ancestor to human in ape G4s; (**B**) from the human-chimpanzee ancestor to chimpanzee in ape G4s; (**C**) from the human-chimpanzee ancestor to human in Hominini G4s; (**D**) from the human-chimpanzee ancestor to chimpanzee in Hominini G4s; (**E**) from the human-chimpanzee ancestor to human in human-specific G4s; (**F**) from the human-chimpanzee ancestor to chimpanzee in chimpanzee-specific G4s; (**G**) from the human-chimpanzee ancestor to human in G4s lost in human; (**H**) from the human-chimpanzee ancestor to chimpanzee in G4s lost in chimpanzee. The substitutions in A, B, C, and D represent changes that do not result in the birth or loss of G4s. Some substitutions in E and F contributed to the birth of G4s. Some substitutions in G and H contributed to the loss (or death) of G4s. For G4s lost in chimpanzee, we used stem and loop annotations of orthologous human G4s. For G4s lost in human, we used stem and loop annotations of orthologous chimpanzee G4s.

For **ape G4s**, the human-chimpanzee ancestor was inferred to have G4s at the same genomic locations as human and chimpanzee. Substitutions in the human or chimpanzee lineages after their divergence were assumed not to result in the birth or loss of G4 motifs. In the human lineage, 22,420 substitutions were identified, while 21,473 substitutions were found in the chimpanzee lineage. The overall substitution frequency at ape G4s was 2.03×10^-3^ substitutions/bp in the human lineage and 2.01×10^-3^ substitutions/bp in the chimpanzee lineage.

More substitutions were found in loops than in stems (Fig. 1B). Correcting for loop and stem lengths, the substitution frequencies at stems and loops were 3.79×10^-4^ substitutions/bp and 3.47×10^-3^ substitutions/bp, respectively, in the human lineage, and 3.54×10^-4^ and 3.45×10^-3^, respectively, in the chimpanzee lineage (Table 1). Therefore, when G4s were maintained in all three species, we observed an approximately one order of magnitude higher substitution frequency in loops than in stems. At ape G4s, substitutions towards G (A->G, C->G, T->G) were common in the stems (Fig. 2A-B), whereas transitions away from Gs were common in the loops (Fig. 2A-B).

**Table 1.** Substitution numbers and frequencies at G4 stems and loops across human and chimpanzee lineages. For G4s lost in chimpanzee, we used stem and loop annotations of orthologous human G4s. For G4s lost in human, we used stem and loop annotations of orthologous chimpanzee G4s.

| Group of G4 | Lineage of substitution | Number of substitution s in stems | Number of substitution s in loops | Lengths of stems (bp) | Lengths of loops (bp) | Substitution frequencies in stems (substitutions/bp) | Substitution frequencies in loops (substitutions/bp) |
| --- | --- | --- | --- | --- | --- | --- | --- |
| Ape | Human | 1,949 | 20,471 | 5,143,374 | 5,893,589 | 0.00038 | 0.0035 |
| Ape | Chimpanzee | 1,768 | 19,705 | 4,990,074 | 5,713,285 | 0.00035 | 0.0034 |
| Hominini | Human | 1,048 | 10,821 | 2,769,897 | 3,185,424 | 0.00038 | 0.0034 |
| Hominini | Chimpanzee | 951 | 10,333 | 2,683,073 | 3,080,198 | 0.00035 | 0.0034 |
| Human-specific | Human | 7,828 | 6,612 | 1,219,254 | 1,390,518 | 0.0064 | 0.0048 |
| Chimpanzee-specific | Chimpanzee | 7,546 | 6,260 | 1,068,349 | 1,221,797 | 0.0071 | 0.0051 |
| G4s lost in chimpanzee | Chimpanzee | 21,850 | 7,872 | 542,803 | 637,664 | 0.040 | 0.012 |
| G4s lost in human | Human | 21,472 | 7,581 | 521,289 | 610,386 | 0.041 | 0.012 |

For **Hominini G4s**, 11,829 substitutions were identified in the human lineage and 11,284 were found in the chimpanzee lineage, resulting in substitution frequencies of 1.99×10^-3^ substitutions/bp and 1.96×10^-3^ substitutions/bp, respectively. Again, more substitutions were found in loops than in stems (Fig. 1B). The substitution frequencies at stems and loops were, respectively, 3.78×10^-4^ and 3.40×10^-3^ in the human lineage, and 3.54×10^-4^ and 3.35×10^-3^ in the chimpanzee lineage (Table 1). This pattern is similar to that observed for ape G4s, suggesting that substitutions that do not lead to birth/loss of G4s are more likely to be found in loops than stems. The substitution spectra at Hominini G4s were similar to those at ape G4s (Fig. 2C-D).

For **human-specific G4s** (Fig. 2E), some nucleotide substitutions from the human-chimpanzee ancestor to the human lineage were responsible for the birth of G4 motifs. We identified a total of 14,440 substitutions at human-specific G4s. Notably, substitutions leading to the birth of G4 motifs occurred in both stem and loop regions (Fig. 1B). Correcting for lengths, the substitution frequencies were 4.76×10^-3^ and 6.42×10^-3^ substitutions/bp in loops and stems, respectively. In the stems of human-specific G4s, substitutions leading to guanine predominated, and transitions (A>G) were more common than transversions (C>G and T>G; Fig. 2E). In contrast, in the loops of human-specific G4s, substitutions away from guanines, particularly G>A transitions, were more frequent than other types of substitutions.

For **chimpanzee-specific G4s**, 6,260 substitutions were found in loop regions, and 7,546 were found in stem regions (Fig. 2B). Correcting for lengths, the frequencies were 5.12×10^-3^ and 7.06×10^-3^ substitutions/bp in loops and stems, respectively (Table 1). The substitution spectrum for chimpanzee-specific G4s was similar to that for human-specific G4s.

For **G4s lost in human**, most substitutions in the human lineage were found in regions corresponding to chimpanzee stems (a total of 21,472), and fewer in regions corresponding to chimpanzee loops (a total of 7,581; Fig. 1B), resulting in substitution frequencies of 4.2×10^-2^ and 1.2×10^-2^, respectively (Table 1). The majority of substitutions originated from guanine (Fig. 2G). Some of these substitutions led to the loss of G4 motifs and, as expected, they occurred in stem regions, highlighting the critical role of stems in maintaining G4 motifs. Similarly, for **G4s lost in chimpanzees**, most substitutions in the chimpanzee lineage were found in regions corresponding to human stems (a total of 21,850) with fewer substitutions occurring in human loops (a total of 7,872; Fig. 1B), resulting in substitution frequencies of 0.040 and 0.012, respectively (Table 1). These substitutions contributed to the loss of G4 motifs in the chimpanzee lineage (Fig. 1), with most originating from guanine (Fig. 2H).

Comparing substitution frequencies between stems and loops of ape G4s shared by multiple species (by chimpanzee and human, or by all three species analyzed; Table 1), we found that loops had substitution frequencies ∼9–10 fold higher than stems (chi-square test *p*-values < 10^-16^, Table S2). In species-specific G4s, stems had substitution frequencies ∼1.4 fold higher than loops (chi-square test *p*-value < 10^-16^, Table S2). In the lineages where G4s were lost, stems had substitution frequencies ∼3-fold higher than loops. Comparing substitution frequencies among evolutionary groups of G4s, we found that G4s shared by multiple species were the most conserved, i.e., had lower substitution frequencies in both stem and loop regions, than newly gained or lost G4s. The highest substitution frequencies were observed in G4s lost in a species-specific manner, suggesting the loss of constraint.

### Stem-loop test for G4s shared by multiple species

We next studied the ratio of substitutions in stems vs. loops in more detail. Relative divergence rates between stems and loops (*D_s_/D_l_*) were calculated as the ratio between substitution frequencies of stems and loops. Since stem regions are more critical to G4 structure formation than loop regions, *D_s_/D_l_* lower than one was considered to be suggestive of G4 motif selection, forming the foundation of a ‘stem-loop test’ of selection for G4s. *D_s_/D_l_* genome-wide was 0.257 in ape G4s (Fig. 3A) and 0.278 in Hominini G4s (Fig. 3B). The similarity in *D_s_/D_l_* ratios between ape and Hominini G4s suggests that both types of G4s experience comparable selective pressures, with a trend to conserve stem regions across evolutionary time. *D_s_/D_l_* at G4s located in non-functional non-repetitive regions (**NFNR**; see Methods) were 0.241 and 0.238 for ape G4s and Hominini G4s, respectively, and can be used as a neutral baseline. For ape G4s (Fig. 3A), promoters, 5’UTRs, and 3’UTRs had particularly low *D_s_/D_l_*values of 0.227, 0.231, and 0.220, respectively, which were significantly lower than the counterparts in NFNR (non-overlapping 95% confidence intervals). For Hominini G4s, we note a particularly low *D_s_/D_l_* = 0.178 at 5’UTRs compared to the counterpart in NFNR (Fig. 3B, Table S4), suggesting particularly strong functional constraints; however, this difference was not statistically significant (overlapping 95% confidence intervals).

**Fig. 3.**
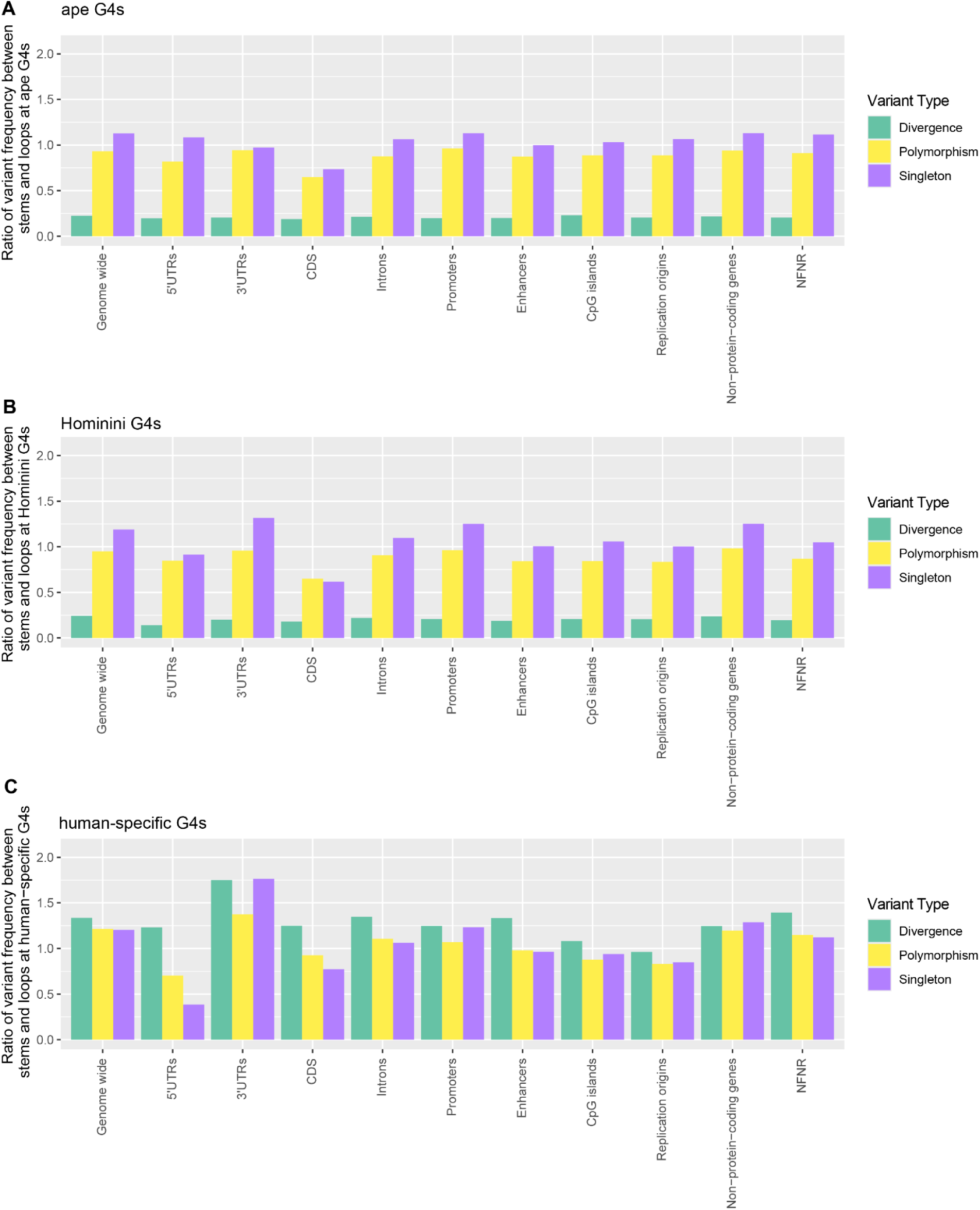
Comparative analysis of ratios of variant frequencies at stems and loops across genomic regions. The ratio of variant frequencies for nucleotide substitutions (i.e., divergence), SNPs (i.e., polymorphism), and singleton SNPs (i.e., singletons) between stems and loops at (**A**) ape G4s; (**B**) Hominini G4s; (**C**) human-specific G4s.

Substitution frequencies depend on both local mutation rates and selective constraints, among other factors. To further understand which of these two factors plays a dominant role in variation in substitution frequency at G4s, we obtained single-nucleotide polymorphisms (SNPs) from the Human Pangenome Reference Consortium (HPRC) dataset (Liao et al.,(Liao et al. 2023). SNPs are expected to be less affected by selection than nucleotide substitutions and to reflect mutation biases more accurately. We calculated SNP frequencies at stems and loops of G4s separately, and then took their ratio. The SNP (i.e., polymorphism) frequency ratio between stems and loops (*P_s_/P_l_*) genome-wide was 1.066 in ape G4s and 1.092 in Hominini G4s. In terms of singletons (polymorphisms found in one human sample), which are expected to be even less affected by selection as compared to all SNPs, the relative singleton mutation rate between stems and loops (*S_s_/S_l_*) genome-wide was 1.367 in ape G4s and 1.292 in Hominini G4s. Thus, compared to nucleotide substitutions (i.e. divergence), SNPs (i.e. polymorphism) are much more evenly distributed between stems and loops, and singleton SNPs have higher rates at stems than loops, suggesting higher mutation rates at guanines than other nucleotides.

These observations strongly suggest that selection acts against fixed differences at stems rather than at loops, which helps maintain G4 structure formation.

### Stem-loop test for human-specific G4s

Human-specific G4s exhibited a markedly different pattern for stem-to-loop ratios for nucleotide substitutions and SNPs compared to G4s shared by multiple species (Fig. 3C). At least some substitutions in human-specific G4s were assumed to be related to G4 birth, reflecting a process where non-G4-forming sequences in ancestral genomes gained G4-forming potential in humans. Their genome-wide *D_s_/D_l_* was 1.65, and their *P_s_/P_l_*was 1.39, indicating that more substitutions were fixed in stems than in loops during G4 birth. The odds ratio of a polymorphism being fixed in loops vs. in stems at human-specific G4s was 0.84.

The relative singleton mutation rate between stems and loops (*S_s_/S_l_*) genome-wide in human-specific G4s was 1.37, which was similar to these ratios in Hominini (1.29) and ape (1.37) G4s. This similarity suggests that mutation rates were higher at stems than loops in all groups of G4, regardless of whether the G4s were maintained, gained, or lost during evolution. This similarity also validates singleton SNPs as a reasonable proxy for the mutation process. While mutation rates were elevated at stems, the fixation of substitutions reflects the selective pressures that shape G4 formation, particularly in regions critical for gene regulation.

### Evaluating selection at G4s with the HKA test

To examine selective pressures on G4 stems and loops, we used two approaches: the Hudson-Kreitman-Aguadé (**HKA**) test (Hudson, Kreitman, and Aguadé 1987) and PhyloFit (Siepel and Haussler 2004). These approaches allowed us to compare fixation probabilities and evolutionary rates across various functional regions in both Hominini and ape G4s.

The **HKA test** (Hudson, Kreitman, and Aguadé 1987) assesses natural selection by comparing patterns of genetic variation within a species (polymorphism) to fixed differences between species (divergence) across multiple loci. Under the neutral theory of evolution, the ratio of polymorphism (*P*) to divergence (*D*) should be similar for the locus being tested, say locus 1, and a locus used as a neutral benchmark, say locus 2. Deviations from the expectation *P_1_*/*D_1_* = *P_2_*/*D_2_* suggest selection: reduced polymorphism vs. divergence indicates positive selection, whereas increased polymorphism vs. divergence suggests balancing or purifying selection. Balancing selection, which maintains multiple alleles at a locus, is unlikely to act on G4s as it would likely disrupt their structural stability and compromise their functionality. Since positive selection is rare genome-wide, it is also unlikely to act on G4s. Therefore, in the case of G4s, the HKA test is mainly expected to measure purifying selection (odds ratio of *P_1_*/*D_1_*/(*P_2_*/*D_2_*) >1).

We first considered whether stems evolved under stronger purifying selection than loops. Using the HKA test, we computed the polymorphism-to-divergence ratio (*P*/*D* ratio) for stems and separately for loops. Then we computed an odds ratio as the *P*/*D* ratio in stems divided by the *P*/*D* ratio in loops. For each type of region (stem or loop), the *P*/*D* ratio was determined by comparing the number of polymorphisms (*P*) within humans to the number of divergent sites (*D*) between human and chimpanzee. When comparing stems to loops, an odds ratio (*P_s_*/*D_s_*/(*P_l_*/*D_l_*)) greater than one suggests stronger purifying selection at stems. Across the entire genome, for polymorphisms with any allele frequencies (not only singletons), the **HKA test** results reveal that the odds ratio in stems versus loops is 3.94 for Hominini G4s and 4.15 for ape G4s (Fig. 4A), suggesting stronger purifying selection operating on stems than on loops. For singletons, the odds ratio between stems and loops is 4.93 for Hominini G4s and 5.01 for ape G4s (Fig. 4B), also suggesting stronger purifying selection at stems. Region-specific HKA test results (Figs. 4A and B) further support these genome-wide patterns. Compared to genome-wide estimates, some functional regions, such as promoters, show particularly high odds ratios for both Hominini G4s (brown) and ape G4s (pink), both when using all polymorphisms and when using singletons. This indicates high conservation of G4 stems in promoters. In contrast, coding sequences (CDS) display low odds ratios, suggesting CDS can tolerate more variation at stems, which could destabilize G4s. Additionally, Hominini G4s show lower odds ratios than ape G4s across most regions, consistent with a weaker selection due to a shorter evolutionary history.

**Fig. 4.**
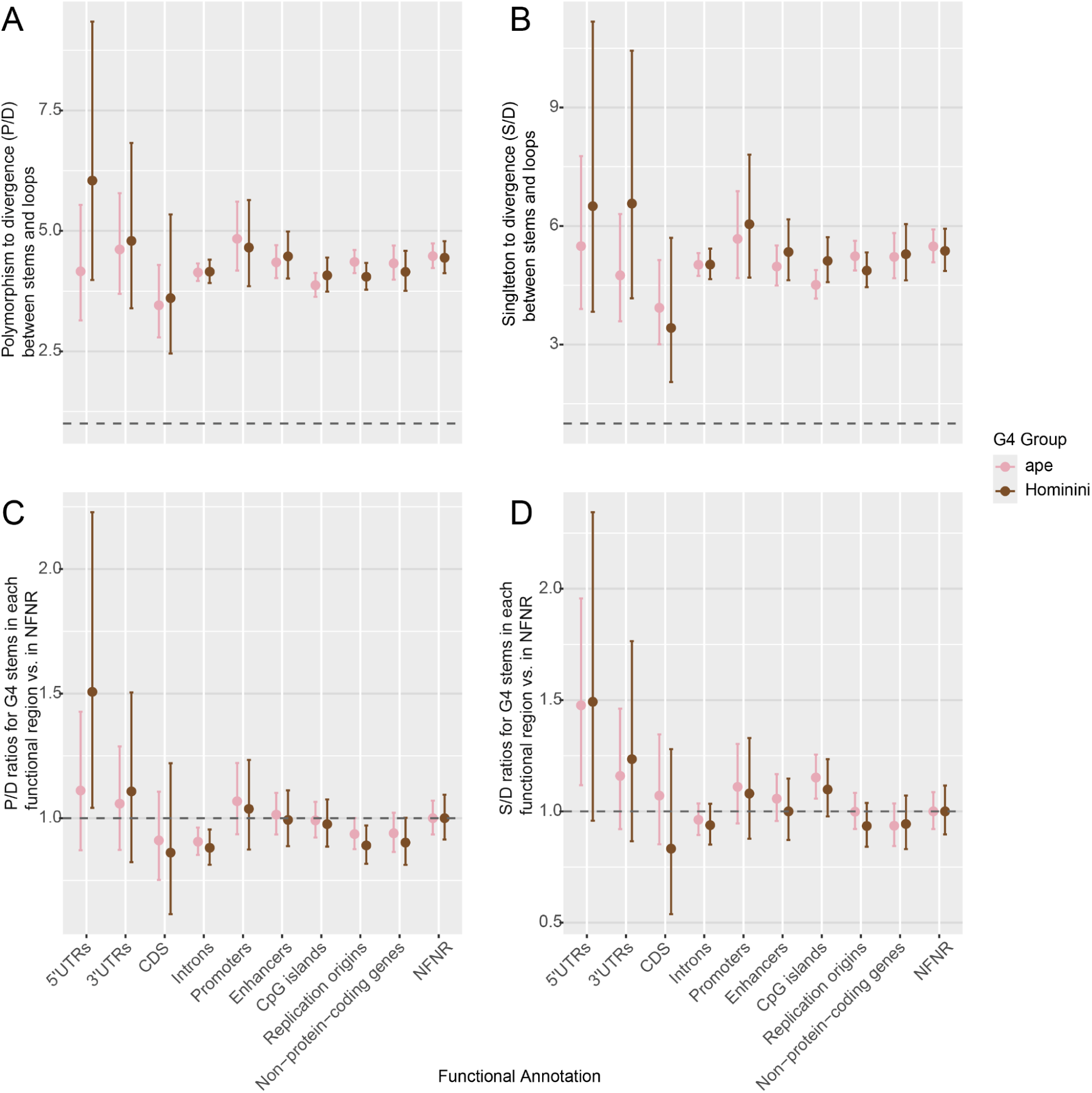
HKA test. Odds ratios (**A**) of polymorphisms to divergence (*P*/*D*) between stems and loops; (**B**) of singletons to divergence (*S*/*D*) between stems and loops; (**C**) of *P*/*D* ratios for whole G4s in each functional region vs. for whole G4 in NFNR (NFNR=1); (**D**) of *S*/*D* ratios for whole G4s in each functional region vs. for whole G4s in NFNR (NFNR=1).

Because our results point to the stem portion of G4s as the main target of selective constraint, we computed odds ratios between the *P*/*D* ratio for G4 stems in different functional regions and *P*/*D* ratio for G4 stems in NFNR. Results obtained using both all polymorphisms and singletons suggest that stems in promoters and UTRs experience markedly stronger purifying selection than stems in NFNR (Fig. 4C–D), whereas stems at CDS show the opposite pattern.

### Measuring selective pressure at G4s with PhyloFit

Last, we applied **PhyloFit** (Siepel and Haussler 2004) to model evolutionary rates between stems and loops across various functional regions in ape and Hominini G4s (Fig. 5A). PhyloFit leverages phylogenetic data to estimate scaling factors based on branch lengths, allowing one to infer lineage-specific evolutionary rates. This approach provides a more precise selection measure by accounting for the evolutionary history of multiple species. Our analysis revealed that the stem-to-loop scaling factor ratios were consistently less than 1 (Fig. 5A), suggesting that, on average, G4 stems evolve more slowly, i.e., likely experience stronger purifying selection, than loops.

**Fig. 5.**
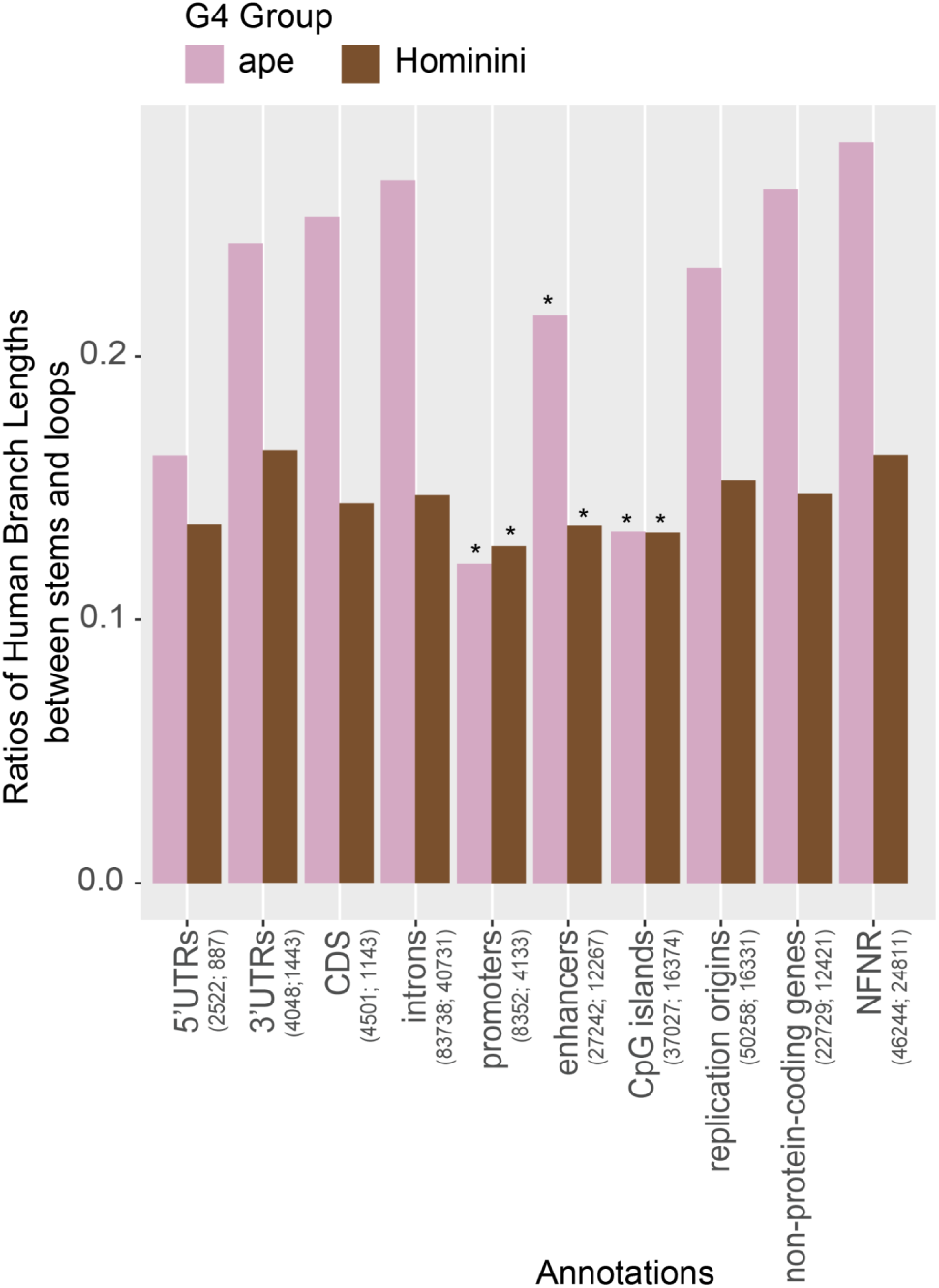
Stem-loop test to compare the selective pressure in stems vs loops. The ratios of the scaling factors between stems and loops in G4s from PhyloFit. X-axis: genomic annotation categories. Numbers in parentheses give the count of overlapping G4s—first for ape G4s, then for Hominini G4s. Asterisks mark permutation-test significance: \**p*-value < 0.05.

The PhyloFit analysis evidenced that regulatory regions exhibit particularly low stem-to-loop scaling factor ratios in both Hominini G4s and ape G4s (Fig. 5A). Specifically, such ratios were significantly lower for G4s in CpG islands, promoters and enhancers than for G4s in NFNR (*p*-values were 0.03, 0.05 and 0.03, respectively, for ape G4s, and 0.04, 0.05 and 0.03, respectively, for Hominini G4s; permutation test, see Methods)–implying stronger purifying selection. This suggests that negative selection acts to preserve G4 motifs in these regions. In contrast, G4 in introns and exons exhibit higher scaling factor ratios than G4s in NFNR; this suggests greater tolerance for variation, with loops showing more flexibility than stems.

For all regions considered but promoters and CpG islands, the ratio of stem-to-loop scaling factors was higher for Hominini than for ape G4s, suggesting less constraint for the former. Overall, Hominini G4s tend to have higher scaling factor ratios than ape G4s (*p*-value=0.03, Wilcoxon test), consistent with weaker selection due to their shorter evolutionary history (i.e., Hominini G4s had less time to accumulate signs of strong selection compared to ape G4s).

## Discussion

Using the recently released T2T great ape genomes, we conducted a comprehensive analysis of substitution dynamics and selection acting on G4 motifs in human, chimpanzee, and Bornean orangutan. Our results showed that most G4 motifs were shared among all three species, some were shared between two species, and only a small subset was species-specific. Substitution dynamics differed between these groups of G4s: in G4s shared by two or more species, changes accumulate mainly in the loops, whereas in species-specific G4s, substitutions predominate in the stems. To evaluate selective pressures and compare selection strength between stems and loops, we applied both the HKA test (Hudson, Kreitman, and Aguadé 1987) and PhyloFit (Siepel and Haussler 2004). We found that stronger purifying selection acted on stems than on loops, particularly in G4s located within promoters, CpG islands, enhancers, and 5′ untranslated regions (5′UTRs).

The conservation of G4 motifs aligns with findings in a recent study (Mohanty, Chiaromonte, and Makova 2025), which demonstrated that G4 motifs evolve according to a molecular clock, maintaining stable rates of evolutionary change across species. A similar trend was also observed in mRNA; 68% of G4s in mRNA were found to be conserved between humans and chimpanzees in terms of motif and structure (Frees et al.,(Frees et al. 2014).

We also found that mutation rates, as inferred from singleton polymorphism frequencies, were higher in stems than in loops across all G4 groups. Prior research (Neupane, Chariker, and Rouchka 2023) also reported higher mutation rates at the start of G4 motifs, which are usually part of the stem region. Among the four types of nucleotides, guanine has the lowest oxidation potential among the DNA bases, making it especially susceptible to oxidative damage (Giorgio et al. 2020). Guanine pairs with cytosine in CpG dinucleotides, which are known to be mutation hotspots (Duncan and Miller 1980). When cytosine in CpG sites is methylated to form 5-methylcytosine, it can spontaneously deaminate to thymine, leading to G:C to A:T transition mutations. Multiple studies showed that G4s are mutation hotspots. The density of nucleotide variants at G4s is higher than in the corresponding control B DNA (Du et al. 2014; Guiblet, Cremona, et al. 2021).

Cross-species comparisons allowed us to unveil additional information. Substitutions that did not lead to the birth or loss of G4 motifs were predominantly found in loops. During G4 birth, most substitutions led to guanines and occurred in stem regions, confirming the essential role of G-rich sequences in forming G4 motifs, and consistent with the earlier conclusion by (Nakken, Rognes, and Hovig 2009). Conversely, when G4 motifs were lost along a phylogenetic branch, most substitutions originated from guanines and also occurred in stems. Previous studies have shown that a single-nucleotide variant from G in stems can lead to the loss of a G4 structure.

For example, a G>A mutation was shown to disrupt a G4 structure in the *c-MYC* promoter, thus increasing the gene’s basal transcriptional activity (Siddiqui-Jain et al. 2002). As another example, single G replacements in the *KRAS* promoter in human and mouse were shown to prevent intramolecular parallel G4 formation, causing the promoter to no longer be repressed by the G4. Mutations in stems can also lead to stabilization/destabilization of G4s in RNA. For example, a G>A mutation in the stem of an RNA G4 at *BCL2* 5’UTR in a lymphoma patient was shown to lower the stability of the G4 and increase mRNA translation efficiency (Zeraati et al. 2017). Conversely, a U>G transversion in the stem of an RNA G4 at *TAOK2* 5’UTR was shown to induce an additional G-tetrad layer, stabilizing a parallel RNA G4 and leading to decreased translation efficiency of mRNA (Zeraati et al. 2017).

Prior research pointed out that mutations in loops or flanking regions may also affect the birth of G4 structures. For example, G4 can be stabilized by introducing mutations to form a hairpin in loops (Nakata et al. 2024). Specifically, a hairpin in the second loop was shown to accelerate the formation of G4s in the presence of KCl *in vitro* (Nakata et al. 2025). Surprisingly, some non-canonical G4 structures containing multiple compensatory mutations in stems can achieve similar or higher biochemical activity including GTP-binding and peroxidase activity (Švehlová et al. 2016).

Both the HKA test and PhyloFit approaches used to evaluate selective pressure consistently revealed stronger purifying selection on stems than loops, particularly in promoters, CpG islands, enhancers, and 5’UTRs. These functional regions are known to be enriched for G4 motifs (Chambers et al. 2015; Guiblet, DeGiorgio, et al. 2021), exhibit hypomethylation (Mohanty, Chiaromonte, and Makova 2025), and contain more stable G4 structures (Guiblet, DeGiorgio, et al. 2021). We propose that the regulatory importance of these regions amplifies the difference in selective pressure between stems and loops, with functional G4s in these regions being under particularly strong purifying selection.

The higher fixation probabilities and PhyloFit scaling factor ratios in Hominini G4s compared to ape G4s suggest weaker selective pressure in the former, likely because of the shorter evolutionary time. These findings suggest that evolutionary time influences the strength of selection on G4 structures, with older ape G4s exhibiting greater conservation and stronger purifying selection, particularly in regions crucial for gene regulation.

While our study advances the understanding of G4 evolution, several limitations should be noted. First, our findings rely on computational predictions using Quadron (Sahakyan et al. 2017), which identifies canonical G4 motifs with G-rich sequences following the pattern G_≥3_N_1-12_G_≥3_N_1-12_G_≥3_N_1-12_G_≥3_. Non-canonical G4s with interrupted G-runs or atypical loop lengths may have been missed, potentially limiting the scope of the analysis. Second, G4s were grouped based on their start and end positions in orthologous sequences, a method that may be overly conservative. This approach could underestimate the number of shared G4s while overestimating lineage-specific G4s. Note that the number of lineage-specific G4s is low in our study despite this potential limitation. Third, our analyses, especially the HKA test, assumed a parsimonious substitution model, which may underestimate the total number of substitutions when reciprocal changes occur at the same site. However, we expect this limitation to be minimal due to the short evolutionary distances considered.

Future research should experimentally validate the predicted G4 structures used in this study to confirm their functional roles and structural stability. Moreover, expanding the analysis to include non-primate species could provide a deeper understanding of the evolutionary origins and conservation of G4s. Finally, experimental approaches such as mutagenesis or CRISPR-based editing could be employed to directly test the impact of stem-specific mutations on G4 stability and function, providing a mechanistic understanding of G4 structures.

## Material & Methods

### Genomic annotations

Human functional annotations were downloaded from https://www.ncbi.nlm.nih.gov/datasets/gene/GCF_009914755.1/. Enhancers were directly extracted from the annotation file. For each of 20,174 protein-coding genes, the longest transcript was selected (using https://gist.github.com/karolpal-jr/48213a0a65475e44f708d5d815127bc3). From the 20,174 transcripts, exons, and CDS were retrieved from the annotation file. Introns were annotated by subtracting exons from mRNAs using bedtools (Quinlan and Hall 2010). 5’UTRs were annotated as the difference in start positions of the 1st exon and the 1st position of the CDS. 3’UTRs were annotated as the difference in end positions of the last exon and the last position in the CDS. Promoters were annotated as 1 kb upstream of genes’ transcription start sites (TSS). Repeat annotations were downloaded from https://hgdownload.soe.ucsc.edu/gbdb/hs1/t2tRepeatMasker/chm13v2.0_rmsk.bb. Unmasked CpG island annotations were downloaded from the UCSC Genome Table Browser for the CHM13v2.0 genome. Non-functional non-repetitive regions (NFNR) were annotated as complements of genes, promoters, enhancers, CpG islands, replication origins, and repeats in the human genome.

### G-quadruplex annotation

G4 loci and their predicted stability were annotated separately in the human, chimpanzee, and B. orangutan genomes using Quadron (Sahakyan et al. 2017). Quadron is a machine-learning-based method trained on a genome-wide dataset generated by G4-seq (Chambers et al. 2015),(Chambers et al. 2015), which experimentally identifies the potential for G4 formation in a genome. The Quadron model computes a score that quantifies the propensity of a DNA sequence to form a stable G-quadruplex structure. The scoring incorporates features from the core G4 motif and the flanking regions, since both regions influence the structural stability of G4s. Higher scores indicate sequences with patterns that facilitate stable folding. The presence and arrangement of guanine bases within and around the motif are particularly influential in determining stability. We segregated the predicted G4 sequences into guanine-rich regions (stems) and the intervening nucleotide sequences (loops) using a regular expression function. The function recognized sequences of three or more consecutive guanine bases as stems, and the sequences separating these stems as loops, which may include guanine bases but cannot consist exclusively of them. The annotated G4 loci were intersected with all the aforementioned functional genomic elements using bedtools (Quinlan and Hall 2010); any overlap of ≥1 bp was considered. To account for the strand-specificity of G4 detection by Quadron, we reverse-complemented the sequence C_3+_N_1−12_C_3+_N_1−12_C_3+_N_1−12_C_3+_ to G_3+_N_1−12_G_3+_N_1−12_G_3+_N_1−12_G_3+,_

### Inferring G-quadruplex presence and sequences in the ancestor

For each great ape species, we extracted the orthologous sequences of G4 loci from the other five great ape species in pruned six-way CACTUS alignments (Paten et al. 2011) using the MAP-SEA pipeline (Mohanty, Chiaromonte, and Makova 2025). G4 loci with the same start and end positions in human, chimpanzee, and B. orangutan, as defined by their whole-genome alignments, were identified as ape G4s (shared by human, chimpanzee, and B. orangutan). G4s only found in human, whose homologous sequences in both chimpanzee and B. orangutan did not contain G4s, were identified as human-specific G4s. G4 loci with the same start and end positions in both human and B. orangutan, but not annotated as G4s in chimpanzee sequences, were identified as G4s lost in the chimpanzee branch. G4 loci with the same start and end positions in both human and chimpanzee, but not annotated as G4s in B. orangutan sequences, were identified as Hominini G4s.

We applied a parsimony-based approach to reconstruct the ancestral genotype and infer the presence or absence of G-quadruplex (G4) structures in the common ancestor ( Table S1). Suppose at a given locus the human sequence matches the orthologous sequence in orangutan (as defined by the CACTUS alignment) but differs from the orthologous sequence in chimpanzee. In that case, we infer that the human-chimpanzee ancestor had the human sequence at that locus. Conversely, suppose the chimpanzee genotype matches the homologous sequence in orangutan but differs from the human sequence. In that case, we infer that the human-chimpanzee ancestor had the chimpanzee sequence at that locus.

### HKA test

To perform the HKA test (Hudson, Kreitman, and Aguadé 1987), we utilized fixed single-nucleotide variants between human and chimpanzee, as well as SNPs and singletons from the human pangenome data (Liao et al. 2023). Fixed and polymorphic mutations were (separately) intersected with functional regions and the NFNR subgenome. We compared the observed counts of fixed and polymorphic variants at G4 stems to those expected based on the loops. We used chi-square test to evaluate the significance of an odds ratio of nucleotide variants to be polymorphic if found at G4 stems vs. loops, as well as the 95% confidence interval of these odds ratios (Fig. 4B). The significance of an odds ratio for singletons was evaluated similarly (Fig. 4A). We also compared the observed counts of fixed and polymorphic variants at G4 stems of a functional region to those expected based on G4 stems at NFNR (Fig. 4D) and applied the same tests to singletons as well (Fig 4C).

### PhyloFit

We used Phylofit to estimate the substitution rates for alignments of each type of G4s. The analysis was performed employing the Expectation Maximization algorithm with medium precision for convergence. We first fit a neutral model (Model A) using alignments of stems from all human G4s located in NFNR regions, and a separate neutral model (Model B) using alignments of loops from all human G4s in NFNR. Then, we fit a new phylogenetic model for the alignments of stems located in different functional regions, using Model A as the background. From this new model, we extracted the scaling factor for stems in each functional region, calculated as the ratio of the branch length for humans between the newly fitted model and the neutral Model A. The same procedure was conducted for loops, using Model B as the background.

We performed a permutation test by randomly sampling two groups of G4s: one group matched the number of G4s in the NFNR set and the other matched the number of G4s in a specific type of regulatory regions (e.g., promoters). For each permutation, we ran PhyloFit to estimate the stem-loop ratios and computed the ratio of these values between the mock NFNR and mock regulatory groups. Repeating this simulation 100 times allowed us to compare the empirical NFNR/regulatory region ratio against the distribution of simulated ratios to assess statistical significance.

## Supplementary Materials

See Supplementary Tables.

## Data Availability

The data underlying this article are available in the article and in its online supplementary material.

## References

Bochman, Matthew L., Katrin Paeschke, and Virginia A. Zakian. 2012. “DNA Secondary Structures: Stability and Function of G-Quadruplex Structures.” Nature Reviews. Genetics 13 (11): 770–80.

Chambers, Vicki S., Giovanni Marsico, Jonathan M. Boutell, Marco Di Antonio, Geoffrey P. Smith, and Shankar Balasubramanian. 2015. “High-Throughput Sequencing of DNA G-Quadruplex Structures in the Human Genome.” Nature Biotechnology 33 (8): 877–81.

Duncan, Bruce K., and Jeffrey H. Miller. 1980. “Mutagenic Deamination of Cytosine Residues in DNA.” Nature 287 (5782): 560–61.

Du, Xiangjun, E. Michael Gertz, Damian Wojtowicz, Dina Zhabinskaya, David Levens, Craig J. Benham, Alejandro A. Schäffer, and Teresa M. Przytycka. 2014. “Potential Non-B DNA Regions in the Human Genome Are Associated with Higher Rates of Nucleotide Mutation and Expression Variation.” Nucleic Acids Research 42 (20): 12367–79.

Frees, Scott, Camille Menendez, Matt Crum, and Paramjeet S. Bagga. 2014. “QGRS-Conserve: A Computational Method for Discovering Evolutionarily Conserved G-Quadruplex Motifs.” Human Genomics 8 (1): 8.

Georgakopoulos-Soares, Ilias, Guillermo E. Parada, Hei Yuen Wong, Ragini Medhi, Giulia Furlan, Roberto Munita, Eric A. Miska, Chun Kit Kwok, and Martin Hemberg. 2022. “Alternative Splicing Modulation by G-Quadruplexes.” Nature Communications 13 (1): 2404.

Giorgio, Marco, Gaetano Ivan Dellino, Valentina Gambino, Niccolo’ Roda, and Pier Giuseppe Pelicci. 2020. “On the Epigenetic Role of Guanosine Oxidation.” Redox Biology 29 (101398): 101398.

Guiblet, Wilfried M., Marzia A. Cremona, Monika Cechova, Robert S. Harris, Iva Kejnovská, Eduard Kejnovsky, Kristin Eckert, Francesca Chiaromonte, and Kateryna D. Makova. 2018. “Long-Read Sequencing Technology Indicates Genome-Wide Effects of Non-B DNA on Polymerization Speed and Error Rate.” Genome Research 28 (12): 1767–78.

Guiblet, Wilfried M., Marzia A. Cremona, Robert S. Harris, Di Chen, Kristin A. Eckert, Francesca Chiaromonte, Yi-Fei Huang, and Kateryna D. Makova. 2021. “Non-B DNA: A Major Contributor to Small- and Large-Scale Variation in Nucleotide Substitution Frequencies across the Genome.” Nucleic Acids Research 49 (3): 1497–1516.

Guiblet, Wilfried M., Michael DeGiorgio, Xiaoheng Cheng, Francesca Chiaromonte, Kristin A. Eckert, Yi-Fei Huang, and Kateryna D. Makova. 2021. “Selection and Thermostability Suggest G-Quadruplexes Are Novel Functional Elements of the Human Genome.” Genome Research 31 (7): 1136–49.

Hubisz, Melissa J., Katherine S. Pollard, and Adam Siepel. 2011. “PHAST and RPHAST: Phylogenetic Analysis with Space/time Models.” Briefings in Bioinformatics 12 (1): 41–51.

Hudson, R. R., M. Kreitman, and M. Aguadé. 1987. “A Test of Neutral Molecular Evolution Based on Nucleotide Data.” Genetics 116 (1): 153–59.

Liao, Wen-Wei, Mobin Asri, Jana Ebler, Daniel Doerr, Marina Haukness, Glenn Hickey, Shuangjia Lu, et al. 2023. “A Draft Human Pangenome Reference.” Nature 617 (7960): 312–24.

Makova, Kateryna D., Brandon D. Pickett, Robert S. Harris, Gabrielle A. Hartley, Monika Cechova, Karol Pal, Sergey Nurk, et al. 2024. “The Complete Sequence and Comparative Analysis of Ape Sex Chromosomes.” Nature 630 (8016): 401–11.

Michael Keesey, T. n.d. “PhyloPic.” PhyloPic. Accessed May 15, 2025. https://www.phylopic.org.

Miga, Karen H., Sergey Koren, Arang Rhie, Mitchell R. Vollger, Ariel Gershman, Andrey Bzikadze, Shelise Brooks, et al. 2020. “Telomere-to-Telomere Assembly of a Complete Human X Chromosome.” Nature 585 (7823): 79–84.

Mohanty, Saswat K., Francesca Chiaromonte, and Kateryna D. Makova. 2025. “Evolutionary Dynamics of Predicted G-Quadruplexes in Human and Other Great Apes.” Genome Biology 26 (1): 161.

Nakata, Minori, Naoki Kosaka, Keiko Kawauchi, and Daisuke Miyoshi. 2024. “Quantitative Effects of the Loop Region on Topology, Thermodynamics, and Cation Binding of DNA G-Quadruplexes.” ACS Omega 9 (32): 35028–36.

Nakata, Minori, Naoki Kosaka, Keiko Kawauchi, and Daisuke Miyoshi. 2025. “Roles of Loop Region in Folding Kinetics and Transcription Inhibition of DNA G-Quadruplexes.” Biochemistry 64 (3): 609–19.

Nakken, S., Torbjørn Rognes, and E. Hovig. 2009. “The Disruptive Positions in Human G-Quadruplex Motifs Are Less Polymorphic and More Conserved than Their Neutral Counterparts.” Nucleic Acids Research 37 (July):5749–56.

Neupane, Aryan, Julia H. Chariker, and Eric C. Rouchka. 2023. “Analysis of Nucleotide Variations in Human G-Quadruplex Forming Regions Associated with Disease States.” bioRxiv. 10.1101/2023.01.30.526341.

Nurk, Sergey, Sergey Koren, Arang Rhie, Mikko Rautiainen, Andrey V. Bzikadze, Alla Mikheenko, Mitchell R. Vollger, et al. 2022. “The Complete Sequence of a Human Genome.” Science 376 (6588): 44–53.

Paten, Benedict, Dent Earl, Ngan Nguyen, Mark Diekhans, Daniel Zerbino, and David Haussler. 2011. “Cactus: Algorithms for Genome Multiple Sequence Alignment.” Genome Research 21 (9): 1512–28.

Quinlan, Aaron R., and Ira M. Hall. 2010. “BEDTools: A Flexible Suite of Utilities for Comparing Genomic Features.” Bioinformatics 26 (6): 841–42.

Rhie, Arang, Sergey Nurk, Monika Cechova, Savannah J. Hoyt, Dylan J. Taylor, Nicolas Altemose, Paul W. Hook, et al. 2023. “The Complete Sequence of a Human Y Chromosome.” Nature 621 (7978): 344–54.

Sahakyan, Aleksandr B., Vicki S. Chambers, Giovanni Marsico, Tobias Santner, Marco Di Antonio, and Shankar Balasubramanian. 2017. “Machine Learning Model for Sequence-Driven DNA G-Quadruplex Formation.” Scientific Reports 7 (1): 1–11.

Sen, D., and W. Gilbert. 1988. “Formation of Parallel Four-Stranded Complexes by Guanine-Rich Motifs in DNA and Its Implications for Meiosis.” Nature 334 (6180): 364–66.

Siddiqui-Jain, Adam, Cory L. Grand, David J. Bearss, and Laurence H. Hurley. 2002. “Direct Evidence for a G-Quadruplex in a Promoter Region and Its Targeting with a Small Molecule to Repress c-MYC Transcription.” Proceedings of the National Academy of Sciences of the United States of America 99 (18): 11593–98.

Siepel, Adam, and David Haussler. 2004. “Phylogenetic Estimation of Context-Dependent Substitution Rates by Maximum Likelihood.” Molecular Biology and Evolution 21 (3): 468–88.

Spiegel, Jochen, Santosh Adhikari, and Shankar Balasubramanian. 2020. “The Structure and Function of DNA G-Quadruplexes.” Trends in Chemistry 2 (2): 123–36.

Sundquist, W. I., and A. Klug. 1989. “Telomeric DNA Dimerizes by Formation of Guanine Tetrads between Hairpin Loops.” Nature 342 (6251): 825–29.

Švehlová, Kateřina, Michael S. Lawrence, Lucie Bednárová, and Edward A. Curtis. 2016. “Altered Biochemical Specificity of G-Quadruplexes with Mutated Tetrads.” Nucleic Acids Research 44 (22): 10789–803.

Varshney, Dhaval, Jochen Spiegel, Katherine Zyner, David Tannahill, and Shankar Balasubramanian. 2020. “The Regulation and Functions of DNA and RNA G-Quadruplexes.” Nature Reviews. Molecular Cell Biology 21 (8): 459–74.

Wang, Guliang, and Karen M. Vasquez. 2023. “Dynamic Alternative DNA Structures in Biology and Disease.” Nature Reviews. Genetics 24 (4): 211–34.

Yoo, Dongahn, Arang Rhie, Prajna Hebbar, Francesca Antonacci, Glennis A. Logsdon, Steven J. Solar, Dmitry Antipov, et al. 2024. “Complete Sequencing of Ape Genomes.” *bioRxiv.org: The Preprint Server for Biology*, October. 10.1101/2024.07.31.605654.

Zeraati, Mahdi, Aaron L. Moye, Jason W. H. Wong, Dilmi Perera, Mark J. Cowley, Daniel U. Christ, Tracy M. Bryan, and Marcel E. Dinger. 2017. “Cancer-Associated Noncoding Mutations Affect RNA G-Quadruplex-Mediated Regulation of Gene Expression.” Scientific Reports 7 (1). 10.1038/s41598-017-00739-y.

Zhang, Zhenjiang, Jixun Dai, Elizabeth Veliath, Roger A. Jones, and Danzhou Yang. 2010. “Structure of a Two-G-Tetrad Intramolecular G-Quadruplex Formed by a Variant Human Telomeric Sequence in K+ Solution: Insights into the Interconversion of Human Telomeric G-Quadruplex Structures.” Nucleic Acids Research 38 (3): 1009–21.

