## Supplementary Table 1-4 for "Substitution spectrum and selection at G-quadruplexes in great ape telomere-to-telomere genomes"

**Table S1.** Pipeline to classify G4s. The 1st column represents source of sequence for each species. The 2nd-7th column shows how to classify G4s. Ape G4s (second column; n=320,000) are G4s annotated by Quadron in human sequence, and also in orthologous sequences of chimpanzee and B. orangutan. G4s lost in chimpanzee (3rd column; n=33,763) shows G4s should be annotated as G4s in both human and the orthologous sequence of B. orangutan, but not annotated in the orthologous sequence of chimpanzee. Hominini G4s (4th column; n=170,012) are annotated as G4s in both human genome and the orthologous sequence of chimpanzee, but not annotated in the orthologous sequence of B. orangutan. Human-specific G4s (5th column; n=72,408) are G4s only annotated in human sequences, but not in the orthologous sequence of B.orangutan and chimpanzee. G4s lost in human (6th column; n=33,552) are G4s annotated in both chimpanzee and the orthologous sequence of B. orangutan, but not annotated in the orthologos sequence of human. Chimpanzee-specific G4s (7th column; n=68,178) are G4s annotated in chimpanzee genome, but not in the orthologous sequence of human and B. orangutan.

| | Ape G4s | G4s lost in chimpanzee | Hominini G4s | Human-specific G4s | G4s lost in human | Chimpanzee-specific G4s |
|---|---|---|---|---|---|---|
| human | Annotated as G4 | Annotated as G4 | Annotated as G4 | Annotated as G4 | Not annotated | Not annotated |
| orthologous sequence of chimpanzee | Annotated as G4 | Not annotated | Annotated as G4 | Not annotated | Annotated as G4 | Annotated as G4 |
| orthologous sequence of B. orangutan | Annotated as G4 | Annotated as G4 | Not annotated | Not annotated | Annotated as G4 | Not annotated |
| Inferred presence of G4s in human-chimpanzee ancestor | Inferred present | Inferred present | Inferred present | Inferred absent | Inferred present | Inferred absent |
| G4 counts | 320,000 | 33,763 | 170,012 | 72,408 | 33,552 | 68,178 |

**Table S2.** Substitution frequency in stems and loops of G4s. The first column identifies the evolutionary lineage in which each substitution occurs, and the second column specifies the G4 group. All subsequent columns present the relevant metrics for each G4 category, conditioned on the substitutions observed in the corresponding lineage.

| Lineage of substitution | Group of G4 | Number of substitutions in stems | Number of substitutions in loops | Lengths of stems (bp) | Lengths of loops (bp) | Substitution frequencies in stems | Substitution frequencies in loops | Number of substi | Length of G4s | overall substitution fre | chi-sqare value | *p*-value |
|---|---|---|---|---|---|---|---|---|---|---|---|---|
| Human | Ape G4s | 1,949 | 20,471 | 5,143,374 | 5,893,589 | 0.00038 | 0.0035 | 22,420 | 11,036,963 | 2.03E-03 | 1.29E+04 | 0 |
| Chimpanzee | Ape G4s | 1,768 | 19,705 | 4,990,074 | 5,713,285 | 0.00035 | 0.0034 | 21,473 | 10,703,359 | 2.01E-03 | 1.27E+04 | 0 |
| Human | Hominini G4s | 1,048 | 10,821 | 2,769,897 | 3,185,424 | 0.00038 | 0.0034 | 11,869 | 5,955,321 | 1.99E-03 | 6.76E+03 | 0 |
| Chimpanzee | Hominini G4s | 951 | 10,333 | 2,683,073 | 3,080,198 | 0.00035 | 0.0034 | 11,284 | 5,763,271 | 1.96E-03 | 6.58E+03 | 0 |
| Human | Human Specific G4s | 7,828 | 6,612 | 1,219,254 | 1,390,518 | 0.0064 | 0.0048 | 14,440 | 2,609,772 | 5.53E-03 | 3.23E+02 | 2.56E-72 |
| Chimpanzee | Chimp Specific G4s | 7,546 | 6,260 | 1,068,349 | 1,221,797 | 0.0071 | 0.0051 | 13,806 | 2,290,146 | 6.03E-03 | 3.53E+02 | 8.51E-79 |
| Chimpanzee | G4s lost in chimpanzee | 21,850 | 7,872 | 542,803 | 637,664 | 0.04 | 0.012 | 29,722 | 1,180,467 | 2.52E-02 | 8.83E+03 | 0 |
| Human | G4s lost in human | 21,472 | 7,581 | 521,289 | 610,386 | 0.041 | 0.012 | 29,053 | 1,131,675 | 2.57E-02 | 8.82E+03 | 0 |

**Table S3.** HKA test to evaluate selection in G4s in genomic regions. ds: number of divergence divided by length in stems. dl: number of divergence divided by length in loops. ps: number of polymorphism divided by length in stems. pl: number of polymorphism divided by length in loops. ss: number of singletons divided by length in stems. sl: number of singletons divided by length in loops.

| G4 group | annotation | G4 counts | Length of functional annotation | stem_length | loop_length | stem_polymorphism | loop_polymorphism | stem_singleton | loop_singleton | stem_divergence | loop_divergence | ds | dl | ps | pl | ss | sl | Ds/Dl | Ps/Pl | Ss/Sl | G4_divergence | G4_polymorphism | G4_singleton |
|---|---|---|---|---|---|---|---|---|---|---|---|---|---|---|---|---|---|---|---|---|---|---|---|
| hominini | CDS | 1141 | 35197319 | 19712 | 26776 | 117 | 180 | 42 | 68 | 54 | 300 | 2.74E-03 | 1.12E | 5.94E- | 6.72E | 2.13E | 2.54E | 2.45E-01 | 8.83E-01 | 8.39E-01 | 354 | 297 | 110 |
| ape | CDS | 4501 | 35197319 | 75209 | 98488 | 379 | 586 | 167 | 227 | 167 | 893 | 2.22E-03 | 9.07E | 5.04E- | 5.95E | 2.22E | 2.30E | 2.45E-01 | 8.47E-01 | 9.63E-01 | 1060 | 965 | 394 |
| human-specific | CDS | 338 | 35197319 | 5932 | 8182 | 63 | 68 | 17 | 22 | 165 | 132 | 2.78E-02 | 1.61E | 1.06E- | 8.31E | 2.87E | 2.69E | 1.72E+00 | 1.28E+00 | 1.07E+00 | 297 | 131 | 39 |
| hominini | CpG islands | 16374 | 43007100 | 297213 | 358430 | 2523 | 2996 | 1055 | 998 | 1028 | 4977 | 3.46E-03 | 1.39E | 8.49E- | 8.36E | 3.55E | 2.78E | 2.49E-01 | 1.02E+00 | 1.27E+00 | 6005 | 5519 | 2053 |
| ape | CpG islands | 37027 | 43007100 | 667812 | 754284 | 4720 | 5324 | 2056 | 1991 | 1911 | 8346 | 2.86E-03 | 1.11E | 7.07E- | 7.06E | 3.08E | 2.64E | 2.59E-01 | 1.00E+00 | 1.17E+00 | 10257 | 10044 | 4047 |
| human-specific | CpG islands | 7116 | 43007100 | 131983 | 161054 | 1790 | 2037 | 586 | 624 | 3459 | 3198 | 2.62E-02 | 1.99E | 1.36E- | 1.26E | 4.44E | 3.87E | 1.32E+00 | 1.07E+00 | 1.15E+00 | 6657 | 3827 | 1210 |
| hominini | Replication origins | 26331 | 107315863 | 457603 | 572780 | 3776 | 4529 | 1472 | 1467 | 1686 | 8186 | 3.68E-03 | 1.43E | 8.25E- | 7.91E | 3.22E | 2.56E | 2.58E-01 | 1.04E+00 | 1.26E+00 | 9872 | 8305 | 2939 |
| ape | Replication origins | 50258 | 107315863 | 845442 | 1015201 | 6058 | 6829 | 2422 | 2272 | 2597 | 12755 | 3.07E-03 | 1.26E | 7.17E- | 6.73E | 2.86E | 2.24E | 2.44E-01 | 1.07E+00 | 1.28E+00 | 15352 | 12887 | 4694 |
| human-specific | Replication origins | 11737 | 107315863 | 220027 | 289293 | 3122 | 3757 | 971 | 1145 | 6405 | 6656 | 2.91E-02 | 2.30E | 1.42E- | 1.30E | 4.41E | 3.96E | 1.27E+00 | 1.09E+00 | 1.12E+00 | 13061 | 6879 | 2116 |
| hominini | Enhancer | 12267 | 55359339 | 203881 | 250581 | 1603 | 1906 | 600 | 597 | 642 | 3414 | 3.15E-03 | 1.36E | 7.86E- | 7.61E | 2.94E | 2.38E | 2.31E-01 | 1.03E+00 | 1.24E+00 | 4056 | 3509 | 1197 |
| ape | Enhancer | 27242 | 55359339 | 449420 | 525330 | 3114 | 3572 | 1216 | 1219 | 1232 | 6145 | 2.74E-03 | 1.17E | 6.93E- | 6.80E | 2.71E | 2.32E | 2.34E-01 | 1.02E+00 | 1.17E+00 | 7377 | 6686 | 2435 |
| human-specific | Enhancer | 4841 | 55359339 | 81331 | 99692 | 964 | 984 | 290 | 301 | 2363 | 1772 | 2.91E-02 | 1.78E | 1.19E- | 9.87E | 3.57E | 3.02E | 1.63E+00 | 1.20E+00 | 1.18E+00 | 4135 | 1948 | 591 |
| hominini | Non-protein-coding genes | 12421 | 488519091 | 200392 | 233932 | 1894 | 1928 | 736 | 588 | 835 | 3528 | 4.17E-03 | 1.51E | 9.45E- | 8.24E | 3.67E | 2.51E | 2.76E-01 | 1.15E+00 | 1.46E+00 | 4363 | 3822 | 1324 |
| ape | Non-protein-coding genes | 22729 | 488519091 | 361599 | 420501 | 2833 | 3023 | 1057 | 935 | 1210 | 5587 | 3.35E-03 | 1.33E | 7.83E- | 7.19E | 2.92E | 2.22E | 2.52E-01 | 1.09E+00 | 1.31E+00 | 6797 | 5856 | 1992 |
| human-specific | Non-protein-coding genes | 5175 | 488519091 | 84096 | 98111 | 1298 | 1086 | 376 | 292 | 2756 | 2213 | 3.28E-02 | 2.26E | 1.54E- | 1.11E- | 4.47E | 2.98E | 1.45E+00 | 1.39E+00 | 1.50E+00 | 4969 | 2384 | 668 |
| hominini | Intron | 40731 | 1128613249 | 663103 | 785462 | 5330 | 5876 | 2109 | 1922 | 2406 | 11017 | 3.63E-03 | 1.40E | 8.04E- | 7.48E | 3.18E | 2.45E | 2.59E-01 | 1.07E+00 | 1.30E+00 | 13423 | 11206 | 4031 |
| ape | Intron | 83738 | 1128613249 | 1351455 | 1575395 | 9407 | 10742 | 3746 | 3526 | 4168 | 19691 | 3.08E-03 | 1.25E | 6.96E- | 6.82E | 2.77E | 2.24E | 2.47E-01 | 1.02E+00 | 1.24E+00 | 23859 | 20149 | 7272 |
| human-specific | Intron | 15998 | 1128613249 | 262135 | 311920 | 3572 | 3229 | 1083 | 1020 | 8042 | 5965 | 3.07E-02 | 1.91E | 1.36E- | 1.04E | 4.13E | 3.27E | 1.60E+00 | 1.32E+00 | 1.26E+00 | 14007 | 6801 | 2103 |
| hominini | NGNR | 24811 | 499600788 | 386345 | 471417 | 3417 | 3939 | 1270 | 1211 | 1359 | 6960 | 3.52E-03 | 1.48E | 8.84E- | 8.36E | 3.29E | 2.57E | 2.38E-01 | 1.06E+00 | 1.28E+00 | 8319 | 7356 | 2481 |
| ape | NGNR | 46244 | 499600788 | 718051 | 852211 | 5923 | 6503 | 2220 | 1990 | 2377 | 11685 | 3.31E-03 | 1.37E | 8.25E- | 7.63E | 3.09E | 2.34E | 2.41E-01 | 1.08E+00 | 1.32E+00 | 14062 | 12426 | 4210 |
| human-specific | NGNR | 10150 | 499600788 | 158367 | 194786 | 2300 | 2001 | 673 | 600 | 5322 | 3820 | 3.36E-02 | 1.96E | 1.45E- | 1.03E | 4.25E | 3.08E | 1.71E+00 | 1.41E+00 | 1.38E+00 | 9142 | 4301 | 1273 |
| hominini | Promoter | 4133 | 80673441 | 67666 | 79523 | 566 | 588 | 219 | 175 | 217 | 1050 | 3.21E-03 | 1.32E | 8.36E- | 7.39E | 3.24E | 2.20E | 2.43E-01 | 1.13E+00 | 1.47E+00 | 1267 | 1154 | 394 |
| ape | Promoter | 8352 | 80673441 | 135659 | 155412 | 942 | 979 | 367 | 325 | 354 | 1780 | 2.61E-03 | 1.15E | 6.94E- | 6.30E | 2.71E | 2.09E | 2.28E-01 | 1.10E+00 | 1.29E+00 | 2134 | 1921 | 692 |
| human-specific | Promoter | 1136 | 19234458 | 19636 | 22770 | 201 | 188 | 74 | 60 | 466 | 373 | 2.37E-02 | 1.64E | 1.02E- | 8.26E | 3.77E | 2.64E | 1.45E+00 | 1.24E+00 | 1.43E+00 | 839 | 389 | 134 |
| hominini | 3'UTR | 1443 | 33909015 | 22753 | 25616 | 181 | 189 | 75 | 57 | 65 | 326 | 2.86E-03 | 1.27E | 7.95E- | 7.38E | 3.30E | 2.23E | 2.24E-01 | 1.08E+00 | 1.48E+00 | 391 | 370 | 132 |
| ape | 3'UTR | 4048 | 33909015 | 65564 | 70757 | 414 | 439 | 170 | 175 | 157 | 769 | 2.39E-03 | 1.09E | 6.31E- | 6.20E | 2.59E | 2.47E | 2.20E-01 | 1.02E+00 | 1.05E+00 | 926 | 853 | 345 |
| human-specific | 3'UTR | 521 | 33909015 | 7807 | 8969 | 99 | 72 | 30 | 17 | 224 | 128 | 2.87E-02 | 1.43E | 1.27E- | 8.03E | 3.84E | 1.90E | 2.01E+00 | 1.58E+00 | 2.03E+00 | 352 | 171 | 47 |
| hominini | 5'UTR | 887 | 6367088 | 15497 | 19670 | 144 | 170 | 53 | 58 | 38 | 272 | 2.45E-03 | 1.38E | 9.29E- | 8.64E | 3.42E | 2.95E | 1.77E-01 | 1.08E+00 | 1.16E+00 | 310 | 314 | 111 |
| ape | 5'UTR | 2522 | 6367088 | 43822 | 51397 | 263 | 321 | 131 | 121 | 95 | 483 | 2.17E-03 | 9.40E | 6.00E- | 6.25E | 2.99E | 2.35E | 2.31E-01 | 9.61E-01 | 1.27E+00 | 578 | 584 | 252 |
| human-specific | 5'UTR | 317 | 6367088 | 5310 | 6818 | 45 | 64 | 10 | 26 | 138 | 112 | 2.60E-02 | 1.64E | 8.47E- | 9.39E | 1.88E | 3.81E | 1.58E+00 | 9.03E-01 | 4.94E-01 | 250 | 109 | 36 |
| hominini | genome wide | 170,012 | 5,955,321 | 2,769,897 | 3,185,424 | 25905 | 27278 | 9976 | 8392 | 11691 | 48439 | 0.0042207 | 0.015 | 0.0093 | 0.008 | 0.003 | 0.002 | 0.2775620 | 1.092130912 | 1.367082089 | 53183 | 60130 | 18368 |
| ape | genome wide | 320,000 | 11,036,963 | 5,143,374 | 5,893,589 | 40988 | 44050 | 15894 | 14096 | 17457 | 77893 | 0.0033940 | 0.013 | 0.0079 | 0.0074 | 0.003 | 0.002 | 0.2568046 | 1.06620952 | 1.292019471 | 85038 | 95350 | 29990 |
| human-specific | genome wide | 72,408 | 2,609,772 | 1,219,254 | 1,390,518 | 19681 | 16188 | 5823 | 4837 | 39226 | 29366 | 0.0321721 | 0.021 | 0.0161 | 0.0116 | 0.004 | 0.003 | 1.5233919 | 1.386552735 | 1.372944965 | 35869 | 68592 | 10660 |

**Table S4.** Stem loop test using HKA test. For every G4 group (second column) that intersects a particular functional annotation (third column) we carried out three separate comparisons—polymorphism, divergence and singletons. In each comparison the odds ratio is defined as (number of variants in stems/total stem length)/(number of variants in loops/total loop length). To test the null hypothesis that this odds ratio equals 1, we constructed a 2 × 2 contingency table of variant versus all sites in stems and loops, then applied a two-sided $\chi^2$ test with one degree of freedom; the resulting $\chi^2$ statistic is converted to a P value via the $\chi^2$ distribution. 95% confidence intervals were shown in last two columns.

| test label | G4 group | annotation | *p*-value | odds ratio | ci_lower | ci_upper |
|---|---|---|---|---|---|---|
| divergence | hominini | CDS | 1.11E-27 | 0.24 | 0.18 | 0.33 |
| | hominini | CpG islands | 0.00E+00 | 0.25 | 0.23 | 0.27 |
| | hominini | Replication origins | 0.00E+00 | 0.26 | 0.24 | 0.27 |
| | hominini | Enhancer | 0.00E+00 | 0.23 | 0.21 | 0.25 |
| | hominini | Non-protein-coding genes | 6.17E-303 | 0.28 | 0.26 | 0.30 |
| | hominini | Intron | 0.00E+00 | 0.26 | 0.25 | 0.27 |
| | hominini | NFNR | 0.00E+00 | 0.24 | 0.22 | 0.25 |
| | hominini | Promoter | 3.10E-103 | 0.24 | 0.21 | 0.28 |
| | hominini | 3'UTR | 2.32E-36 | 0.22 | 0.17 | 0.29 |
| | hominini | 5'UTR | 2.04E-33 | 0.18 | 0.12 | 0.25 |
| | hominini | genome wide | 0.00E+00 | 0.28 | 0.27 | 0.28 |
| | ape | CDS | 1.14E-81 | 0.24 | 0.21 | 0.29 |
| | ape | CpG islands | 0.00E+00 | 0.26 | 0.25 | 0.27 |
| | ape | Replication origins | 0.00E+00 | 0.24 | 0.23 | 0.26 |
| | ape | Enhancer | 0.00E+00 | 0.23 | 0.22 | 0.25 |
| | ape | Non-protein-coding genes | 0.00E+00 | 0.25 | 0.24 | 0.27 |
| | ape | Intron | 0.00E+00 | 0.25 | 0.24 | 0.26 |
| | ape | NFNR | 0.00E+00 | 0.24 | 0.23 | 0.25 |
| | ape | Promoter | 9.08E-187 | 0.23 | 0.20 | 0.26 |
| | ape | 3'UTR | 3.12E-87 | 0.22 | 0.18 | 0.26 |
| | ape | 5'UTR | 9.72E-51 | 0.23 | 0.18 | 0.29 |
| | ape | genome wide | 0.00E+00 | 0.26 | 0.25 | 0.26 |
| | human-specific | CDS | 4.34E-06 | 1.72 | 1.36 | 2.19 |
| | human-specific | CpG islands | 4.77E-29 | 1.32 | 1.26 | 1.39 |
| | human-specific | Replication origins | 5.29E-40 | 1.27 | 1.22 | 1.31 |
| | human-specific | Enhancer | 1.42E-54 | 1.63 | 1.54 | 1.74 |
| | human-specific | Non-protein-coding genes | 2.32E-38 | 1.45 | 1.37 | 1.54 |
| | human-specific | Intron | 1.14E-166 | 1.60 | 1.55 | 1.66 |
| | human-specific | NFNR | 4.05E-141 | 1.71 | 1.64 | 1.79 |
| | human-specific | Promoter | 1.22E-07 | 1.45 | 1.26 | 1.67 |
| | human-specific | 3'UTR | 1.96E-10 | 2.01 | 1.61 | 2.52 |
| | human-specific | 5'UTR | 3.83E-04 | 1.58 | 1.22 | 2.05 |
| | human-specific | genome wide | 0.00E+00 | 1.52 | 1.50 | 1.55 |

| | | | | | | |
|---|---|---|---|---|---|---|
| | hominini | CDS | 3.17E-01 | 0.88 | 0.69 | 1.12 |
| | hominini | CpG islands | 5.69E-01 | 1.02 | 0.96 | 1.07 |
| | hominini | Replication origins | 5.51E-02 | 1.04 | 1.00 | 1.09 |
| | hominini | Enhancer | 3.32E-01 | 1.03 | 0.97 | 1.11 |
| | hominini | Non-protein-coding genes | 2.61E-05 | 1.15 | 1.08 | 1.22 |
| | hominini | Intron | 1.58E-04 | 1.07 | 1.04 | 1.12 |
| | hominini | NFNR | 1.58E-02 | 1.06 | 1.01 | 1.11 |
| | hominini | Promoter | 3.79E-02 | 1.13 | 1.01 | 1.27 |
| | hominini | 3'UTR | 4.97E-01 | 1.08 | 0.87 | 1.33 |
| | hominini | 5'UTR | 5.30E-01 | 1.08 | 0.85 | 1.35 |
| | hominini | genome wide | 5.27E-24 | 1.09 | 1.07 | 1.11 |
| | ape | CDS | 1.21E-02 | 0.85 | 0.74 | 0.97 |
| | ape | CpG islands | 9.52E-01 | 1.00 | 0.96 | 1.04 |
| | ape | Replication origins | 3.71E-04 | 1.07 | 1.03 | 1.10 |
| | ape | Enhancer | 4.45E-01 | 1.02 | 0.97 | 1.07 |
| | ape | Non-protein-coding genes | 1.10E-03 | 1.09 | 1.03 | 1.15 |
| | ape | Intron | 1.46E-01 | 1.02 | 0.99 | 1.05 |
| | ape | NFNR | 1.62E-05 | 1.08 | 1.04 | 1.12 |
| | ape | Promoter | 3.48E-02 | 1.10 | 1.01 | 1.21 |
| | ape | 3'UTR | 8.10E-01 | 1.02 | 0.89 | 1.17 |
| | ape | 5'UTR | 6.47E-01 | 0.96 | 0.81 | 1.14 |
| | ape | genome wide | 1.41E-20 | 1.07 | 1.05 | 1.08 |
| | human-specific | CDS | 1.82E-01 | 1.28 | 0.89 | 1.83 |
| | human-specific | CpG islands | 3.35E-02 | 1.07 | 1.01 | 1.14 |
| | human-specific | Replication origins | 2.87E-04 | 1.09 | 1.04 | 1.15 |
| | human-specific | Enhancer | 6.11E-05 | 1.20 | 1.10 | 1.31 |
| | human-specific | Non-protein-coding genes | 9.11E-16 | 1.39 | 1.28 | 1.51 |
| | human-specific | Intron | 2.19E-29 | 1.32 | 1.25 | 1.38 |
| | human-specific | NFNR | 1.89E-29 | 1.41 | 1.33 | 1.50 |
| | human-specific | Promoter | 3.62E-02 | 1.24 | 1.01 | 1.52 |
| | human-specific | 3'UTR | 3.34E-03 | 1.58 | 1.15 | 2.17 |
| | human-specific | 5'UTR | 6.29E-01 | 0.90 | 0.60 | 1.35 |
| polymorphism | human-specific | genome wide | 9.09E-207 | 1.39 | 1.36 | 1.42 |

| | | | | | | |
|---|---|---|---|---|---|---|
| singletons | hominini | CDS | 3.86E-01 | 0.84 | 0.56 | 1.25 |
| | hominini | CpG islands | 4.04E-08 | 1.27 | 1.17 | 1.39 |
| | hominini | Replication origins | 7.34E-10 | 1.26 | 1.17 | 1.35 |
| | hominini | Enhancer | 2.73E-04 | 1.24 | 1.10 | 1.39 |
| | hominini | Non-protein-coding genes | 6.65E-12 | 1.46 | 1.31 | 1.63 |
| | hominini | Intron | 9.97E-17 | 1.30 | 1.22 | 1.38 |
| | hominini | NFNR | 8.88E-10 | 1.28 | 1.18 | 1.39 |
| | hominini | Promoter | 1.43E-04 | 1.47 | 1.20 | 1.80 |
| | hominini | 3'UTR | 2.87E-02 | 1.48 | 1.04 | 2.13 |
| | hominini | 5'UTR | 4.45E-01 | 1.16 | 0.78 | 1.71 |
| | hominini | genome wide | 4.70E-99 | 1.37 | 1.33 | 1.41 |
| | ape | CDS | 7.22E-01 | 0.96 | 0.78 | 1.18 |
| | ape | CpG islands | 1.09E-06 | 1.17 | 1.10 | 1.24 |
| | ape | Replication origins | 3.34E-17 | 1.28 | 1.21 | 1.36 |
| | ape | Enhancer | 1.65E-04 | 1.17 | 1.08 | 1.26 |
| | ape | Non-protein-coding genes | 1.18E-09 | 1.31 | 1.20 | 1.44 |
| | ape | Intron | 9.12E-20 | 1.24 | 1.18 | 1.30 |
| | ape | NFNR | 1.13E-19 | 1.32 | 1.25 | 1.41 |
| | ape | Promoter | 7.78E-04 | 1.29 | 1.11 | 1.51 |
| | ape | 3'UTR | 6.66E-01 | 1.05 | 0.84 | 1.30 |
| | ape | 5'UTR | 6.63E-02 | 1.27 | 0.98 | 1.64 |
| | ape | genome wide | 1.30E-108 | 1.29 | 1.26 | 1.32 |
| | human-specific | CDS | 8.72E-01 | 1.07 | 0.53 | 2.10 |
| | human-specific | CpG islands | 1.90E-02 | 1.15 | 1.02 | 1.29 |
| | human-specific | Replication origins | 1.30E-02 | 1.12 | 1.02 | 1.22 |
| | human-specific | Enhancer | 4.67E-02 | 1.18 | 1.00 | 1.39 |
| | human-specific | Non-protein-coding genes | 1.75E-07 | 1.50 | 1.29 | 1.76 |
| | human-specific | Intron | 9.60E-08 | 1.26 | 1.16 | 1.38 |
| | human-specific | NFNR | 1.11E-08 | 1.38 | 1.23 | 1.54 |
| | human-specific | Promoter | 4.56E-02 | 1.43 | 1.00 | 2.05 |
| | human-specific | 3'UTR | 1.91E-02 | 2.03 | 1.08 | 3.92 |
| | human-specific | 5'UTR | 6.36E-02 | 0.49 | 0.21 | 1.06 |
| | human-specific | genome wide | 9.21E-60 | 1.37 | 1.32 | 1.43 |
